# Mapping the structure of drosophilid behavior

**DOI:** 10.1101/002873

**Authors:** Gordon J. Berman, Daniel M. Choi, William Bialek, Joshua W. Shaevitz

## Abstract

Most animals possess the ability to actuate a vast diversity of movements, ostensibly constrained only by morphology and physics. In practice, however, a frequent assumption in behavioral science is that most of an animal’s activities can be described in terms of a small set of stereotyped motifs. Here we introduce a method for mapping the behavioral space of organisms, relying only upon the underlying structure of postural movement data to organize and classify behaviors. We find that six different drosophilid species each perform a mix of non-stereotyped actions and over one hundred hierarchically-organized, stereotyped behaviors. Moreover, we use this approach to compare these species’ behavioral spaces, systematically identifying subtle behavioral differences between closely-related species.

---

Animals perform a complex array of behaviors, from changes in body posture to vocalizations to other dynamic outputs. Far from being a disordered collection of actions, however, there is thought to be an intrinsic structure to the set of behaviors and their temporal organization [1, 2]. While behavior can be thought of as a trajectory through a large dimensional space, there is evidence in several different systems that natural behaviors do not fill this space uniformly, but rather are confined to lower-dimensional manifolds [3–5]. Moreover, the paths an animal takes through this reduced space are thought to be decomposable into sequences of stereotyped motions or modules [6–8].

Understanding the nature of this type of dimensional reduction is central to discussion of problems ranging from neural coding to evolution [9–13], but the lack of a comprehensive and compelling mathematical framework for behavioral analysis has limited progress. Standard paradigms for the study of behavior often rely on the use of intuitive definitions of behavior and probe their occurrence using methods ranging from human observation to supervised machine-learning algorithms [14–17]. Here we attempt a more systematic approach to the analysis of behavior in six closely related species of flies, developing tools that allow objective definitions of stereotypy, hierarchy, and similarities in behavioral phenotype.

The basis of our approach is to view behavior as a trajectory through a high-dimensional space of postural dynamics. Given some natural coordinate system on this space, stereotyped behaviors are ones in which the trajectory hovers near particular, repeatable positions. Thus the task of behavioral analysis is to start with raw data (here, high resolution movies) and construct this “behavioral space” *B*, spanned by coordinates, 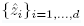 These coordinates must be complete, in that they can represent all observed actions. The probability distribution over *B*, *b*(**z**), measures the relative fractions of time spent on different actions, and peaks in this distribution are candidates for stereotyped behaviors.

Both stereotyped and non-stereotyped behaviors can be exhibited over time as an animal moves within this behavioral space. When we can see well-resolved peaks in the distribution *b*(**z**), and the trajectory pauses in the neighborhood of these peaks, then we can say that we have identified a stereotyped behavior. Non-stereotyped actions, on the other hand, correspond to epochs in which the trajectory moves rapidly through behavioral space without pausing at characteristic locations. These data—driven, unsupervised definitions do not require us to assume, *a priori*, that stereotyped actions must exist. We will also see that the spatial structure of *b*(**z**) provides a framework to explore the relationships between stereotyped actions.

We explore here the spontaneous behaviors of ground-based flies in a largely featureless circular arena [Figs. 1A, 5]. Under these conditions, flies display a multitude of complex, non-aerial behaviors such as locomotion and grooming, typically involving multiple parts of their bodies. To capture dynamic rearrangements of the fly’s posture, we recorded video of individual behaving animals with sufficient spatiotemporal resolution to resolve moving body parts such as the legs, wings, and proboscis. Each animal was imaged at 100 Hz for one hour, yielding 3.6 × 10^5^ movie frames per individual, and in each frame we focus our analysis on a 200 × 200 pixel square containing the fly. In these experiments, we studied six species of *Drosophila*: *D. mauritiana, D. simulans, D. sechellia, D. melanogaster, D. santomea*, and *D. yakuba* (four individuals each, corresponding to 1.44 × 10^6^ images per species and over 8.5 million images in total).

The number of postures an animal can adopt is limited by the mechanical properties of its body. The fly body is made up of relatively inflexible segments connected by mobile joints, so that the number of postural degrees of freedom is relatively small, and it is tempting to try extracting these variables directly from the images, but occlusions and the complex fly geometry make this difficult [18, 19]. As an alternative, we note that almost all of the variance in the 4 × 10^4^ pixel images can be explained by projecting onto a Euclidean space of just 50 dimensions or principal components (Figs 1C and D, Appendix E), after correcting for rigid rotations and translations of the fly’s body; we refer to these dimensions as postural modes. Thus we can convert a movie of fly behavior into a 50—dimensional time series, 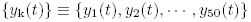 representing the projections onto each of the postural modes over time (Fig 1E).

**FIG. 1.**
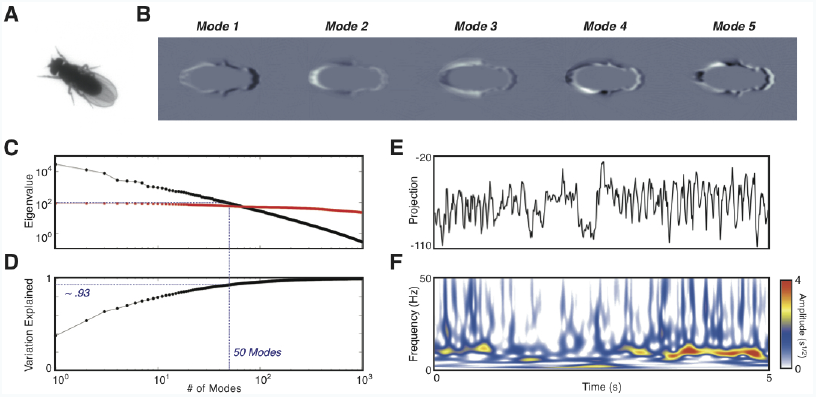
Quantitative description of behavioral movies. (A) Raw image of a fly in the arena. (B) Pictorial representation of the first 5 postural modes, 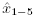, after inverse Radon transform. Black and white regions represent highlighted areas of each mode (with opposite sign). (C) First 1,000 eigenvalues of the data matrix (black) and shuffled data (red). (D) Fraction of cumulative variation explained as a function of number of modes included. (E) Typical time series of the projection along postural mode 6 and (F) its corresponding wavelet transform.

The postural modes themselves, however, do not provide a fully natural coordinate system for behavior. In particular, there is a challenge in recognizing behaviors that correspond to the similar sequences of movements at different times and with different durations [14, 15, 20]. As an alternative, we consider a spectrogram representation, where we measure the power at frequency *f* associated with each mode k, in a window surrounding a moment in time, *S*(k, *f*; *t*) (Fig 1F, Appendix E). In practice we compute the spectrum using Morlet wavelets, and capture dynamics from *f* = 1 to *f* = 50 Hz, corresponding to 25 independent frequency channels. *S*(k, *f*; *t*) is thus a 1,250 dimensional feature vector at each time *t*. We expect that the dynamics do not fill this space, and there are several ways of searching for lower dimensional structures [21]. The approach we use here is to project the features *S*(k, *f*; *t*) onto two dimensions, *z*_1_(*t*) and *z*_2_(*t*), such that we preserve the local neighborhood relations among points but allow for distortions on longer length scales (Appendix H) [22].

Figure 2A shows the embedding of all the spectral feature vectors that we observed, across all six species, into the space *z*_1_, *z*_2_. Although our embedding method is invariant to variations in the total power of the postural motions 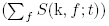 we see that nearby points have similar total power. If we coarse grain this image to generate an estimate of local density *b*(**z**), we see (Fig 2B) there are large number of resolved local maxima that are candidates for stereotyped behaviors. If we look at the dynamics of *z*_1_(*t*) and *z*_2_(*t*), we find that the system really does pause, with near zero velocity, at points corresponding to maxima of the probability density (Fig 2D).

**FIG. 2.**
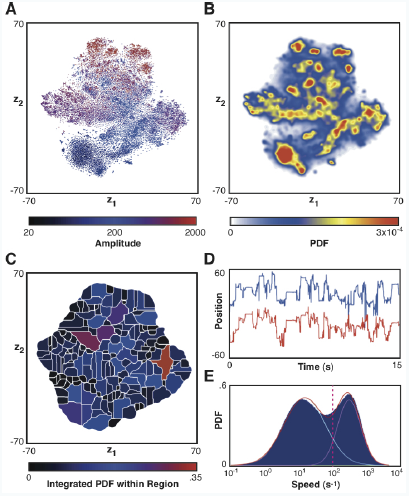
Embedding into behavioral space. (A) Results from embedding spectral feature vectors, {*S*(*k*, *f*; *t*)}, into 2D. Points are color coded by the summed amplitude of the wavelet feature vector. (B) Estimated probability density function (PDF) from embedding all points. (C) Integral of the PDF over each of the discretized regions. (D) Trajectory segment through behavioral space, *z*_1_(*t*) (blue) and *z*_2_(*t*) (red). (E) Histogram of velocities in the embedded space fit to a two-component log-gaussian mixture model. The blue bar chart represents the measured probability distribution, the red line is the fitted model, the cyan and green lines are the mixture components of the fitted model, and the red dashed line is the threshold between the high and low velocity states.

The impression that trajectories pause can be made more precise by looking at the distribution of speeds through the space **z**, 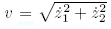. In Fig 2E, we see that the distribution *P*(*v*) is, in fact, bimodal, and that the two peaks are separated by nearly a factor of 100 in speed. It makes sense to think of this distribution as a mixture of two states, “paused” and “moving,” and we can then assign every moment in time a probability of being in the paused state. Pauses occur preferentially at the peaks of the distribution *b*(**z**) (Fig. 18), and with a reasonably conservative threshold we can say that the fly is paused at some point in the behavioral space **z** almost precisely half the time (*f*_pause_ = 0.4997 across all individuals and species studied here, Fig. 15). We identify these pauses near peaks of *b*(**z**) as stereotyped behavioral states or actions: they persist, with residence times in one state range from ∼ .05 s out to more than 10 s; Fig. 15, they recur many times over the one hour observation of an individual, and they they can be identified across multiple individuals.

We can make our impression of multiple peaks in the probability distribution *b*(**z**) precise by asking for connected regions in the *z*_1_, *z*_2_ plane such that climbing up the gradient of probability density always leads to the same local maximum. In image processing this is called a “watershed transform” [23], and if the probability distribution comes from a physical system in thermal equilibrium this is equivalent to finding the valleys in the free energy landscape. We show the results of this analysis in Fig 2C, where we identify 169 distinct regions of the behavioral space, each of which surrounds a single local maximum of probability density. Although this analysis is based on pooled data form all the species, each species visits almost all of these states during the course of our observations, with *D. melanogaster* visiting the fewest (134) and *D. simulans* the most (157); see Fig. 15, Table II. Importantly, visual inspection of the movies in epochs where the behavior sits within one of these distinct regions often allows us to recognize familiar behaviors: walking, running, wing grooming, and even proboscis extension (Fig. 3). These stereotyped behaviors, which correspond to nearly stationary values of *z*_1_ and *z*_2_, correspond to periodic orbits in the space of postural modes (Appendix J, Fig. 24), showing that we have mapped temporal sequences into single points within *B*.

**FIG. 3.**
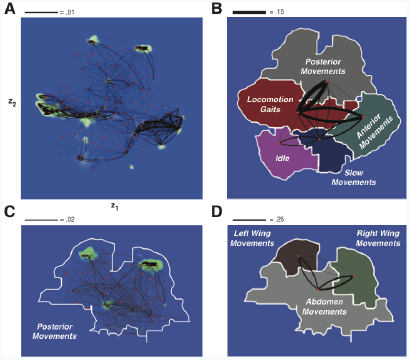
The structure of behavioral space for *D. mauritiana.* (A) Transitions between the 155 watershed regions observed in the species. Each red point represents the maximum of the local PDF, and the black lines represent the transition probabilities between the regions. Black line thicknesses are proportional to the transition flux (ignoring self-transitions), and right-handed curvature implies the direction of transmission. For clarity, all lines representing fluxes less than 10^−4^ are omitted. (B) Behavioral labels associated with each partitioned graph component. Black line thicknesses are proportional to conditional transition probabilities between regions. (C) Zoom-in on the ”Posterior Movements” module. Lines now represent conditional transition probabilities to the next observed state within the module. (D) Further partitioning of the graph from (C) yields more fine-grained details.

Having found discrete states, we can measure the transition probability *T*_*ij*_ that the system will follow its visit to state *i* immediately by a visit to a different state *j* (Appendix I). We created the space **z** by requiring that neighborhood relations be preserved, and correspondingly we find that transitions are more likely among nearby states (Fig 3A). Looking at the physical movements associated with these different states, we see, for instance, that a walking fly is much more likely to transition to turning or walking at a different speed in the next bout of activity than it is to groom its wing, or that a fly that is grooming its eyes is more likely to groom its antennae in the next bout than it is to extend its proboscis. While these seem plausible, it should be emphasized that with more than one hundred discrete states, and thus many thousands of possible transitions, not all the structure is so intuitive.

It is tempting to think that the locality of transitions corresponds to a modular organization of behavior: if the system is in state *i*, it is more likely to transition to states *j* within the same module, while inter—module transitions are more rare. If this is correct, then knowing which module the system is in should capture a large fraction of the information available about the next state that will be visited, and this information theoretic approach leads to an algorithm for partitioning the set of states into components [24, Appendix I]. We find that the resulting graph components are spatial segregated in the **z** plane, and correspond to broad behavioral categories (Fig. 3B). If we allow for more graph components, still trying to capture the most information about the next state, we find that these broad categories break into smaller components, hierarchically (Fig. 3B-D), and this is true for all six species we observed (Fig. 22).

The fact that the different species we have studied can be described in a common behavioral space **z** means that we can use the distribution over this space to compare the species to one another. Thus, we can construct the probability distribution *b*(**z**) separately for each species (Fig 4A), and we can measure the pairwise distances between these distributions using the Jensen-Shannon (JS) divergence, which quantifies how much information one sample of the behavioral state provides about the choice between the two species that might have generated that sample [25, Fig. 4B-E, Eq. K1]. When the species are ordered according to their molecular phylogeny [26, 27], we identify a high degree of behavioral similarity within the most recently evolved clade consisting of *D. mauritiana*, *D. simulans*, and *D. sechellia*. This clade diverged in the past ≈ 250,000 years [28] and we find small within-clade JS divergences compared to the larger JS divergences with the other species. It is less clear, however, that the JS divergence is the proper metric to track evolutionary distance on longer scales.

**FIG. 4.**
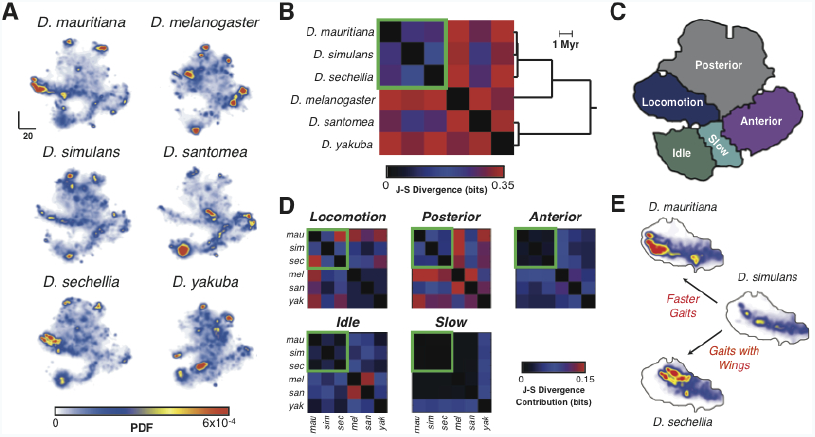
Comparative behavioral spaces. (A) Estimated PDFs for each species individually. (B) Jensen-Shannon divergences between all pairs of species. The emanating tree is compiled from the molecular-based phylogenies found in [26, 27]. The green square highlights flies of the *D. simulans* clade. (C) Partition of behavioral space based upon transition matrices from all species. (D) Contributions of each of the modular regions to the overall J-S divergence (the sum of all matrices is equal to the matrix in (B)). Note that most differences within the highlighted clade are within the ”Locomotion” module. (E) Zoom-in of the locomotion portion of the PDF for the three highlighted species. *mauritiana* is observed to perform higher-frequency gaits than *simulans*, as shown by its shift leftward within the region. *sechellia* displays similar-frequency motion to *simulans*, but often performs wing motions during locomotion gaits, as shown by its upward shift.

To probe the origin of the small behavioral differences seen from species in this clade, we separately calculated the contribution of each major behavioral region (Fig. 4C) to the overall JS divergence (Fig. 4D). Nearly half (49.7%) of the total variation in this clade was due to differences in locomotion behaviors. To a lesser extent (29.2%), posterior behaviors differed, and very little behavioral difference was observed for idle, slow, and anterior behaviors. Comparing locomotory behaviors, we find that *D. mauritiana* and *D. sechellia* are the most distinct, with *D. simulans* being the nearest species to both (Fig. 4B,D). Behaviorally, we see that the predominate contributor to this separation is the existence of higher-frequency locomotion gaits in *D. mauritiana* and the presence of gaits involving wing motions in *D. sechellia* (Figs. 4E, 23). This pattern of differences, with *D. sechellia* and *D. mauritiana* more similar to *D. simulans* than to each other, is consistent with the ecological observation that the former two species are island endemics that likely diverged from *D. simulans*, a cosmopolitan human commensurate [28, 29].

The ability to compare the structure of behavioral space and the organization of stereotyped motions between populations of behaving animals has applications beyond the study of behavioral evolution in fruit flies. Combined with tools for genetic manipulation, DNA sequencing, neural imaging, and electrophysiology, the identification of subtle behavioral distinctions and patterns between groups of individuals will impact deep questions related to the interactions between genes, neurons, behavior, and evolution. In this initial study, we probed the motion of individuals in a largely featureless environment. Extensions to more complicated situations, e.g. where sensory inputs are measured and/or controlled, or multiple individuals are present, are easily implemented. Finally, we note that the only *Drosophila*-specific step in our analysis pipeline is the generation of the postural eigenmodes. Given movies of sufficient quality and length from different organisms, spectral feature vectors and behavioral spaces can be similarly generated, allowing for potential applications from worms to mice to humans and a greater understanding of how animals behave.

## Acknowledgements

We thank Kieren James-Lubin, and Kelsi Lindblad for assistance in data collection and analysis as well as David Stern, David Schwab, and Thibaud Taillefumier for discussions and suggestions. J.W.S. and G.J.B. also acknowledge the Howard Hughes Medical Institute Janelia Farm Visitor Program and the Aspen Center for Physics, where many ideas for this work were formulated. This work was funded through awards from the National Institutes of Health (GM098090, GM071508), The National Science Foundation (PHY–0957573, PHY–1066293), the Pew Charitable Trusts, the Swartz Foundation, and the Alfred P. Sloan Foundation.

## Appendices

### Appendix A: Fly-Tracking Instrument

To acquire high-resolution image data of ground-based flies, we built a tracking instrument that allows us to track a single fly as it moves freely in a large arena (Figure 5). This apparatus consists of a camera and lens system (Gazelle Point Gray, 100 Hz, 1088 × 1088 pixels, bit depth of 8) that streams images directly to disk and a high-speed X-Y translation stage (Griffon Motion) that is used to keep the fly in the field of view. Backlighting is provided by LED illumination and the magnification is chosen such that a single fly is approximately 100 pixels in length and has an area that covers 6,000-8,000 pixels. The entire set-up is encased in an opaque box, preventing external visual stimuli.

**FIG. 5.**
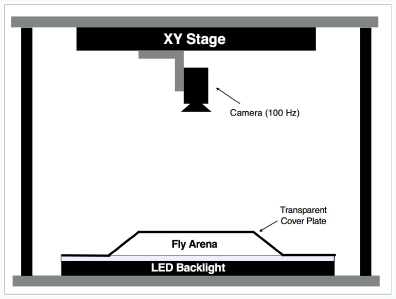
A schematic of the recording apparatus.

We designed our arena based on previous work which showed that a thin chamber with gently sloping sides prevents flies from flying, jumping, and climbing the walls [30]. To keep the behaving flies in the focal plane of our camera, we inverted the previous design. Our arena consists of a vacuum-formed, clear PETG plastic dome 4 inches in diameter with sloping sides at the edge clamped to a flat glass plate. The edges of the plastic cover are sloped to prevent the flies from being occluded and to limit their ability to climb upside-down on the cover. The latter of these aims is further aided by applying Sigma Cote to the underside of the cover, thus preventing adhesion to the surface. In practice, we find that this set-up results in effectively no bouts of upside-down walking.

A Proportional-Integral-Derivative (PID) feedback algorithm is used to keep the moving fly inside the camera frame by controlling the position of the X-Y stage based on the camera image in real time. The feedback parameters are tuned such that the camera tracks a fly smoothly in a sufficiently fast manner to capture all observed terrestrial manoeuvres. All aspects of the instrumentation are controlled by a single computer using a custom-written LabView graphical user interface.

This setup is able to record millions of images without losing the fly from the field of view. A sample trajectory of fly motion derived from the tracking routine is displayed in Figure 6. Due to storage limitations, we only save a 200 × 200 pixel square that is centred on the centroid of the fly. Data used in this paper consists of movies of 24 individual flies from the *Drosophila melanogaster* species subgroup, each recorded for one hour (approximately 9 million frames in total).

**FIG. 6.**
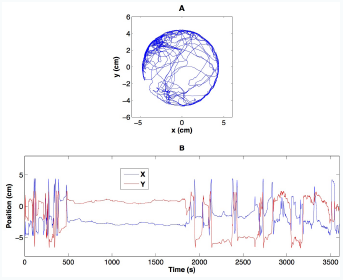
A) Example Time-trace of a fly as it moves about the filming arena. B) X and Y positions plotted individually.

### Appendix B: Animals

All flies used were male from the strains listed in Table I. These flies were isolated and reared separately within 4 hours of ecolsion and data was collected at 12-14 days after eclosion. All recording occurred between the hours of 9:00 AM and 1:00 PM, thus reducing the effect of circadian rhythms. All recordings for an individual species were performed during a single day. Flies were placed into the arena via aspiration and were subsequently allowed 10 minutes for adaptation before data collection. The temperature for all recordings was 25.5° ± .5°*C.*

**Table I.**
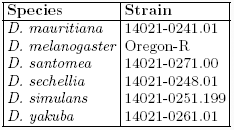
Strains used

| Species | Strain |
| --- | --- |
| *D. mauritiana* | 14021-0241.01 |
| *D. melanogaster* | Oregon-R |
| *D. santomea* | 14021-0271.00 |
| *D. sechellia* | 14021-0248.01 |
| *D. simulans* | 14021-0251.199 |
| *D. yakuba* | 14021-0261.01 |

### Appendix C: Data Analysis

The general framework of our analysis is outlined in Fig. 7 as well as the main text. Images are first segmented and registered. After this, they are decomposed into postural time series and converted into wavelet spectrograms. These spectrograms are used to construct spectral feature vectors that we embed into 2 dimensions using t-Distributed Stochastic Neighbor Embedding [22]. Lastly, we use this embedding to create definitions for behavioral states. Each of these steps are described in the subsequent sections. All calculations are performed using custom-written MATLAB code executed on a Mac Pro desktop computer containing two 2.66 GHz 6-core Intel Xeon processors and 64 GB of RAM.

**FIG. 7.**
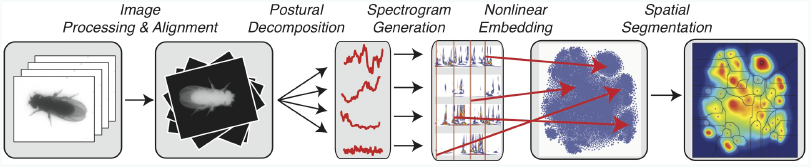
Schematic of the data analysis pipeline.

### Appendix D: Image Processing

Image preprocessing is done in two steps: segmentation and registration.

#### 1. Segmentation

The first phase of our data analysis is to isolate the insect within the original image. First, we invert the image, leading to values near zero for the background and maximal possible values of 255 within the fly. As a first pass, we set all pixels below a set threshold (40) to zero to eliminate small noise in the background. Then we apply Canny’s method for edge detection [31], resulting in a binary image containing the edge positions. We then morphologically dilate this binary image by a 3 × 3 square in order to fill any spurious holes in the edges and proceed to fill all closed curves. This filled image is then morphologically eroded by a square of the same size, resulting in our desired segmentation. If the resulting mask is smaller than a minimal value (i.e. because a hole along the edge is not filled), the size of the dilation and the sensitivity of the edge detection are adjusted until an area threshold is met. For our purposes, we set the area threshold to be 85% of the size of the previous image, and the first image is monitored for accuracy prior to subsequent segmentations.

#### 2. Registration

While our tracking algorithm ensures that the fly remains within the image boundaries, the center of the fly and the orientation within the frame vary over time. Having obtained a sequence of isolated fly images, we next register them both translationally and rotationally with respect to a template image via a method similar to that developed previously [32–34]. The template image is generated by taking a typical image of a fly and then manually ablating the wings and legs digitally (Figure 8, top right).

**FIG. 8.**
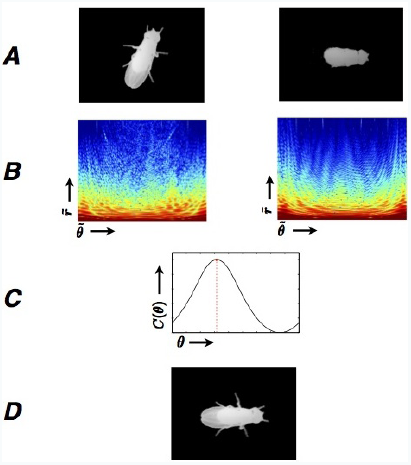
Outline of the alignment algorithm. Starting with the segmented image and a pre-selected template image (A, left and right, respectively), magnitudes of the polar Fourier transforms are computed (B). From these, the phase correlation between the Fourier magnitude angles are computed, and the maximal value of this function is deemed the rotation angle (C). Finally, the original image is rotated through this angle, and the translation in the x-y plane is found in a similar manner, resulting in the aligned image (D).

The rotational alignment is performed by taking advantage of the property that for two images, *I*_1_(*x*, *y*) and *I*_2_(*x*, *y*), that are translated and rotated with respect to each other by 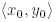 and *θ*_0_, respectively, their 2D Fourier transforms, 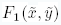 and 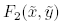 can be related by

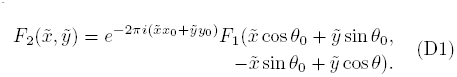

Information about shifts in position can therefore be eliminated by looking only at the magnitude of the spectra, 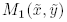, and 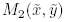. This principle ensures that rotational and translational alignment can be performed independently.

In polar coordinates, we have

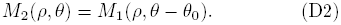

Practically, we compute *M*_1_ and *M*_2_ by taking 1D FFTs along the *r*-axis of the Radon transforms of the original images, with the Radon transform defined as

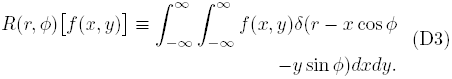

This procedure is equivalent to finding the polar Fourier spectra as a result of the projection-slice theorem, which states that a slice through the 2D Fourier Transform of an image can be calculated by taking the 1D Fourier Transform of a projection onto a line through the same original image [35]. Given *M*_1_ and *M*_2_, *θ*_0_ can be found by maximising the average cross-correlation between the two amplitude images along the *θ* axis. Specifically, this is done by solving the following optimization problem numerically,

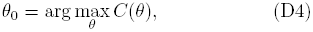

where

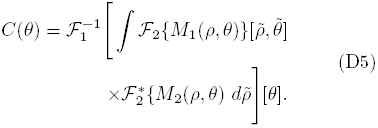

Here, 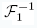 is a inverse 1D Fourier Transform and 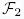 is a 2D Fourier Transform.

The orientations calculated in this way have a 180° degeneracy. We overcome this obstacle by manually assigning the direction of the first frame in a sequence and then assuming that the insect moves less than 180° in each successive frame. After rotating the original image through the desired angle, we rescale the fly so that it’s body – not including wings and legs – has the same numberof pixels as the template image, allowing for effective comparisons between multiple flies. This essentially assumes that fly bodies differ predominately via an overall scalar multiplier. Lastly, translational alignment is achieved through use of a sub-pixel method for finding the offset that maximizes the phase-correlation between the images [36].

### Appendix E: Postural Representation and Dimensional Reduction

#### 1. Subsampling and variance thresholding

The segmented fly images contain too many pixels to analyse using our memory-limited techniques. We therefore seek to represent the segmented and aligned images in a manner that preserves nearly all of the information in the image sequence while at the same time compressing the data. The first operation we perform to curb this excess is to down-sample our segmented and aligned images by a factor of 10/7 after alignment (from 200 × 200 to 140 × 140 pixels). The fly possesses no features that are less than 2 pixels wide even at this reduced resolution. Even after coarse-graining, we do not need to study the entire 19,600-dimensional system. Many pixels within the fly image are either always zero or always saturated and thus contain no dynamical information. Accordingly, we would like to use only a subsample of these measurements.

The most obvious manner to go about this is to find the pixels containing the highest variance and keep only those above a certain threshold. The primary difficulty here, however, is that there is not an obvious truncation point (Figure 9A). This is most likely the result of the fact that the fly legs can potentially occupy the majority of the pixels in the image, but only are present in a relatively small number in any given frame. Hence, many of these periphery pixels all have similar, moderate standard deviations, making them difficult to differentiate as important or negligible.

**FIG. 9.**
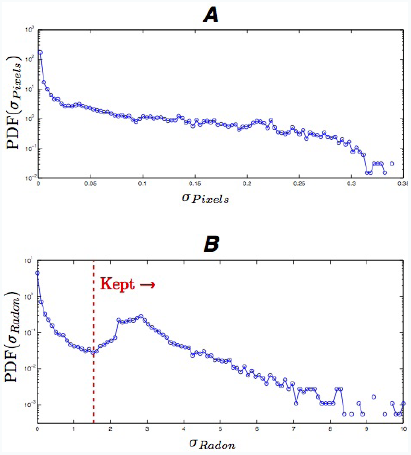
A) Probability density function of pixel standard deviations. B) Probability density function of Radon pixel standard deviations. Note the clear minimum that exists in B), allowing for an effective reduction in the number of pixels necessary to represent the data.

A more compact scheme is to represent the images in Radon-transform space (Equation D3), which more sparsely parameterizes lines such as legs or wing veins. After Radon transformation, the probability density function of pixel-value standard deviations has a clear minimum and we keep pixels whose standard deviation is larger than this value (Figure 9B). This results in keeping 6,763 pixels out of 18,090 which retain approximately 95% of the total variation in the images.

#### 2. Principal Components Analysis of Fly Body Posture

Principal components analysis (PCA) is a frequently-used method for converting a set of correlated variables into a set of values of linearly uncorrelated eigenmodes. PCA also facilitates dimensional reduction when a limited set of components explains a large amount of the total observed variation in the data. Results from this analysis can be described as the space spanned by the eigenvectors of the data covariance matrix, *C*, corresponding to the largest *m* eigenvalues out of the total latent dimensionality of the data, *δ*. While in general there is no rigorous manner to choose *m*, we discuss a way to estimate an upper bound in *m* in Section E3.

As seen in Figure 11, the majority of the observed variation occurs in a relatively small number of modes. If we define variation explained as

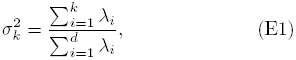

where λ_*i*_ is the *i*th largest eigenvalue of *C*, we find that 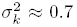 for *k* = 7, 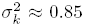 for *k* = 14, and 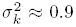 for *k* = 24. These data lack any characteristic kink in the eigenvalue spectrum, the traditional tell-tale sign of where to truncate the representation. This is most likely a result of the fact that the low-dimensional manifold on which the data lies is non-linear, resulting in an effective mixing between lower-variation modes. Figure 12 shows the inverse Radon transform of the first 49 eigenmodes of *C*. The modes are all fly-like and are relatively localized at particular points, usually along the edges of the fly body and wings. While several of these postural modes are localized to discrete appendages on the fly body, in general a one-to-one assignment between the eigenmodes and the fly physiology is not possible. As a result, we shall rely primarily on frequency-domain projections for behavioral analyses.

**FIG. 10.**
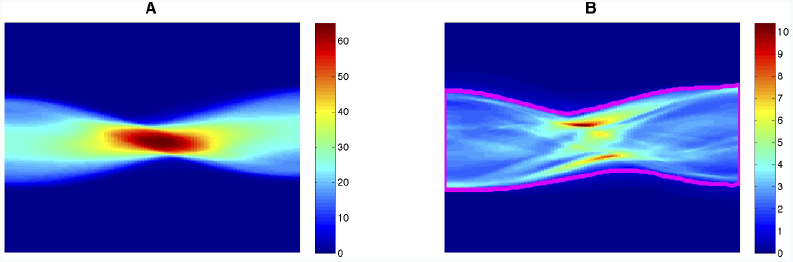
Mean and standard deviations of the Radon-transformed images, blue being zero and red being maximal. **A** shows the mean values, and **B** displays the corresponding standard deviations of the data set. The region bounded by the magenta line in Figure **B** represents the subset of pixels used for further analysis.

**FIG. 11.**
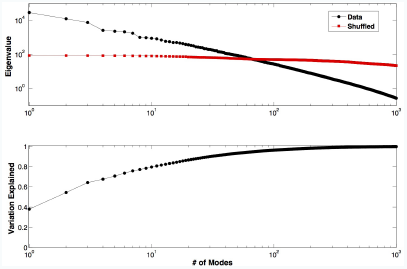
A) Plot of the first 1,000 eigenvalues of C (black circles) and the noise model (red squares). 50 modes from our data PCA are larger than the largest noise eigenvalue. B) The cumulative variance explained in the data.

**FIG. 12.**
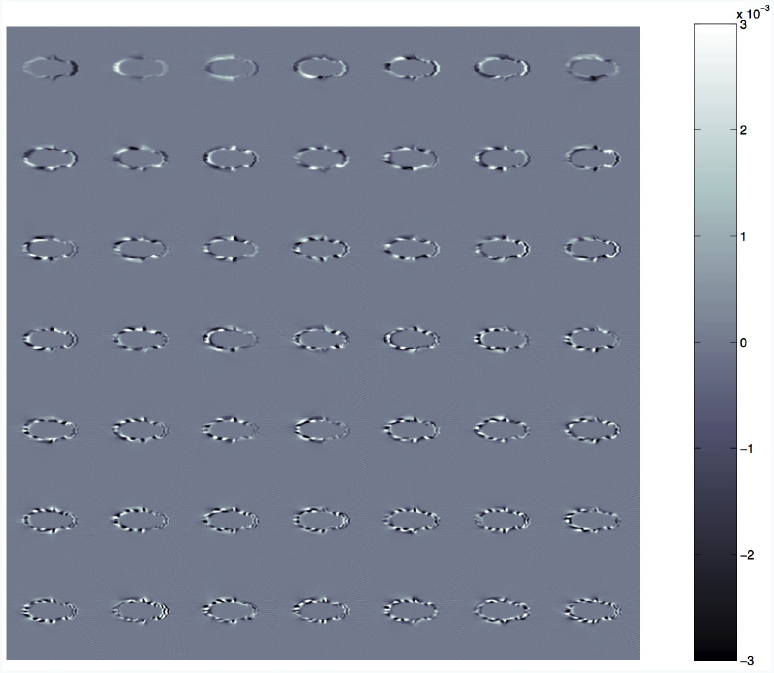
The 49 most significant eigenmodes (after inverse Radon transformation), displayed in descending order (left-to-right then top-to-bottom). The color scale is in arbitrary units.

#### 3. Series truncation and comparison to the null model

Dimensional reduction of postural space via truncation from the PCA is difficult because of the smoothness of the eigenvalue spectrum. Given this, one approach to determining the truncation point is to compare the PCA eigenvalues with a null model based on the noise properties of our data set. This noise is due to finite data collection as well as any measurement error in both the tracking software and image preprocessing and registration. Eigenvalues that are statistically indistinguishable from zero (i.e. modes whose amplitude is below the noise threshold) are not included in the subsequent dynamic analyses. This process is aided by the sharp-cutoffs found in random matrix eigenvalues.

Our specific null model here is that each of the Radon pixel values are drawn independently from each other but out of the same distribution as seen experimentally.

In other words, we take our data matrix, 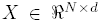(rows corresponding to individual images and columns corresponding to Radon space pixels), and shuffle each of the columns independently from each other. In this model, given an infinite number of samples, there should be no linear correlations. Given finite sampling (even if very large), however, there will still remain some residual correlations, resulting in off-diagonal non-zero terms in the covariance matrix. Hence, if we diagonalize this new covariance matrix, the largest eigenvalue provides a resolution limit for our ability to distinguish signal from finite sampling noise.

Performing this null model analysis, we find that 50 modes from our data set have eigenvalues greater than the largest observed eigenvalue from the null model (Fig. 11, top). These 50 modes account for slightly more than 93% of the observed variation in the data (Fig. 11, bottom). Subsequent dynamic analysis then uses this low-dimensional postural representation, 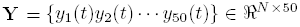, to represent the full image data set.

It should be noted that this procedure places an upper-bound on the number of modes necessary to represent the fly postures. There are undoubtably more sources of noise in our data set than finite sampling error alone (i.e. segmentation and alignment errors), but setting the eigenvalue cut-off as described eliminates modes that are unambiguously indistinguishable from noise.

### Appendix F: Wavelet Analysis and Power Spectrum Calculation

Given a low-dimensional postural representation, **Y**, we wish to discover sets of image sequences that represent stereotyped behaviors. Clustering of the postures, however, clearly does not capture the dynamics of the fly motion. For example, a stationary fly should be classified as “resting” regardless of the positions of its appendages. Similarly, different bouts of a periodic walking gate should be classified together despite their different phases or durations during a measurement.

We attempt to ameliorate these difficulties by analyzing **Y** via a time-frequency analysis. Many time-frequency analyses implement a gaussian-windowed Fourier Transform (sometimes referred to as a Gabor filter or a short-time Fourier transform):

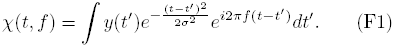

While this results in a smooth, well-behaved output, the uncertainty in the temporal and frequency axes are linked such that 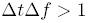. Decreasing the filter width, *σ*, requires a corresponding increase in 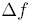. This presents a fundamental difficulty because behavioral events occur over many different time scales, yet we desire the maximum temporal resolution possible.

One way to cope with this fundamental limitation is to retile the sampling of time-frequency space. Namely, we wish to sample low-frequency events over longer time windows, and higher-frequency over shorter time windows, thereby giving instantaneous measures of both long time-scale behaviors and sharp transitions. A convenient formalism for achieving this tiling that maintains accuracy in both the time and frequency dimensions is the continuous wavelet transform (CWT). Although this transform shares many similarities with the Fourier transform, primarily a result of the fact that both can be viewed as rotations in a functional space, wavelets possess the desired multi-resolution time-frequency trade-off and often can represent the data in a more sparse manner [37]. Specifically, wavelets are generated by morphing a mother wavelet, 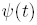, through a scaling and translation,

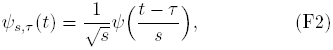

where 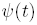 has a mean of zero, is square-normalizable, has a Fourier transform, 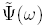, that obeys

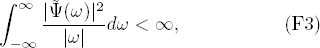

and is otherwise well-behaved. These conditions insure that 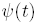 is localized and well-defined for all *t*. We then define a set of wavelet coefficients, 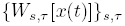, via

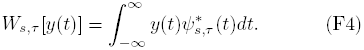

Applying the convolution theorem to Eq. F4 and using the definition of the mother wavelet from Eq. F2, we have

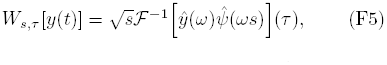

where the hat notation indicates a Fourier transformed variable and 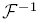 is the inverse Fourier transform. Given an analytical formula for 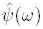, Eq. F5 provides a fast and reliable method for computing the wavelet coefficients.

The fly behavioral data generally consists of long, stationary sequences interrupted by bursts of activity. Morlet’s wavelet function,

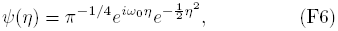

is able to capture these chirps while being relatively insensitive to noise [38]. This function has the additional property that the scale, *s*, is related to the Fourier frequency, *f*, by

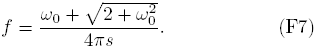

This can be derived by maximizing 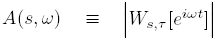 with respect to *s*.

Because of the differing time scales used for each of the wavelet coefficient calculations, this leads to a bias that makes *A(s, ω)* disproportionally large when responding to pure sine waves of lower frequencies. To correct for this, we find a scalar function *C(s)* such that

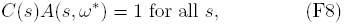

where *ω*\* is 2π times the Fourier frequency found in Eq. F7. For a Morlet wavelet, this function is

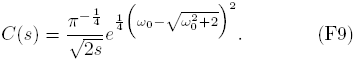

Combining this result with Eq. F5, we can define the power spectrum

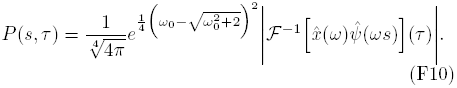

By picking andyadically-spaced set of frequencies between *f*_*min*_ = 1 Hz and the Nyquist frequency (*f*_*max*_ = 50 Hz) via

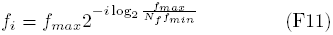

for *i* = 1, 2,…, *N*_*f*_ (and their corresponding scales via Eq. F7), one can create a wavelet spectrogram that is resolved at multiple time-scales. Accordingly, we can build a multimodal wavelet data set by applying Eq. F10 to each of the first 50 modes for *N_f_* = 25 frequencies between 1 and 50 Hz, yielding a matrix 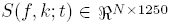.

### Appendix G: Distance Metric and Intrinsic Dimensionality

Starting from the matrix *S*(*f*, *k*; *t*), each row of which contains the wavelet transform centered about a particular moment in time, we would like to possess a distance metric that provides the basis for the space in which we wish to embed our data set. Our choice of metric hinges on two important observations. First, because the data results from the amplitudes of wavelet transforms, all entries in *S*(*f*, *k*; *t*) must be non-negative. Second, as a particular behavior starts, sustains, and ends, the feature vector will change by an overall multiplicative factor, but will vary only slightly in shape. Accordingly, since we possess a set of non-negative feature vectors, we can eliminate this variance by normalizing each vector separately. In other words, we let

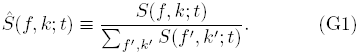

Given that we now have a set of feature vectors that are all non-negative and normalized to one, a natural distance metric between these points can be derived by treating each feature vector as a probability density over all mode-frequency channels. The natural distance metric between two such probability distributions is the Kullback-Leibler divergence between them [25]. Mathematically, this means that we compute the distance between two time points, *t*_1_ and *t*_2_, via

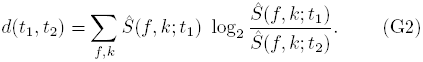

This can be viewed as the extra number of bits required to encode 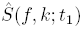 if based on a code built upon 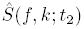 rather than itself. It should be noted at this point that our definition for *d*(*t*_1_, *t*_2_) is not symmetric, but this does not cause a problem for the embedding procedure described in the next section.

Given this distance metric, it is now possible to get a rough estimate of the dimensionality of *M*, the manifold populated by the set of all values for 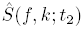, and show that it is much smaller than 1,250. The most straight-forward means for performing this is to calculate the correlation dimension, *d_C_*(*M*) of our data. In this method, we estimate *d*(*M*) via

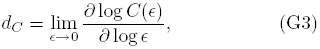

where

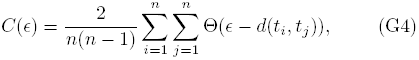

*n* is the number of data points, and 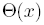 is the Heaviside step function. Thus, by fitting *C*(*∊*) as a function of *∊* on a log-log plot to a line (excluding regions of saturation at high *∊* and under-sampling at low ∊), we can obtain an estimate for *d*_c_ (Fig. 13). Here, we find that *d_C_* ≈ 12.7. While this is only an estimate and by no means a statement of the implicit dimensionality of the manifold containing the data (which can potentially vary from point to point), it confirms that our data has a dimensionality much less than our naive 1,250-dimensional parameterization.

**FIG. 13.**
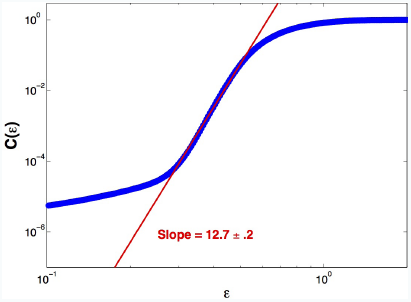
Estimation of the correlation dimension of the data set (Eq. G4). The blue line is *C* as a function of 6, and the red line is a linear fit to the middle region of the plot. The measured slope is 12.7 ± .2.

### Appendix H: t-Distributed Stochastic Neighbor Embedding

This section gives an outline of the t-Distributed Stochastic Neighbor Embedding (t-SNE) algorithm [22] and details as to our implementation of it. Like other embedding algorithms, t-SNE aims to take data from a high-dimensional space and embed it into a space of much smaller dimension, preserving some set of invariants as best as possible.

For t-SNE, the conserved invariants are related to the Markov transition probabilities if a random walk is performed on the data set. Specifically, let us assume that we have *n* time points in our data set, and the transition probability from time point *t_i_* to time point *t_j_*, *P_j|i_*, is proportional to a Gaussian kernel of the distance between them:

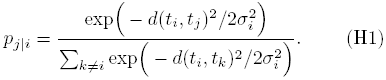

All self-transitions (i.e. *p_i|i_*) are assumed to be zero. The t-SNE algorithm attempts to embed the data such that these transition probabilities are optimally preserved. Each of the *σ_i_* are set such that all points have the same transition entropy, 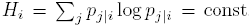, via binary search until |2^*H*_0_^ − 2*^H_i_^* | < 10^−5^. Here, we set 2*^H_i_^* = 30, which can be thought of as a proxy for selecting the number of nearest neighbors to which a point will transition.

We also need to define transition probabilities in the embedded space. The naive approach, initially outlined in [39], is to assume that the transitions in this space, *q_j|i_*, are also given via Gaussian kernels. This becomes problematic, however, because the process of embedding objects of relatively high intrinsic dimension (say, 10), results in a crashing of points towards the centre of the map, thus obscuring the clustering properties of the original space. t-SNE alleviates this problem by mandating that transition probabilities in the embedded space be proportional to a heavy-tailed distribution. Specifically, the Student-t distribution is used:

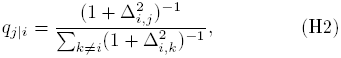

where Δ_*i,j*_ is the Euclidean distance between embedded points 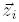 and 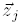, corresponding to *t_i_* and *t_j_*, respectively. Note that no values for *σ* are required in this space, since they would only result in an overall multiplicative constant.

If we symmetrize these transition probabilities by defining

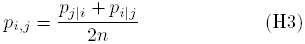

and

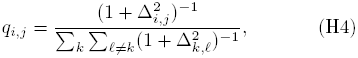

then our cost function to be minimized is the Kullback-Leibler divergence between the *p_i,j_* and *q_i,j_*, or

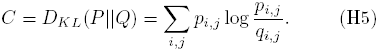

This cost function is used – as opposed to a cost function relying on the unsymmetrized probabilities – due to ease of optimization and previous studies have reported that it results in similar maps as the symmetric version [22, 40]. In general, though, this choice of cost function is beneficial because it places low weight on points that are far apart in the original space (low *p_i j_*), while still maintaining information about the manifold's local structure. This separates it from methods such as diffusion maps [41] or multi-dimensional scaling [42], where effects from far-away points can dominate the cost function.

Unfortunately, Equation H5 is not convex, so some care is necessary to optimize it. Specifically, we optimize this cost function via gradient descent, utilizing techniques proposed in the original t-SNE paper [22]. Given the cost function in Equation H5, it is possible to show that

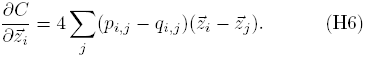

If *χ*^(*n*)^ is the solution of the equations at iteration *n*, we follow the gradient downward using

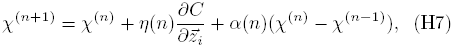

where *η*(*η*) is an adaptive learning rate that is set via the scheme from [43], and *α*(*η*) is a momentum term. Here, we set *η*(1) = 100 and

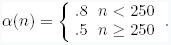

In addition, we utilize the “early exaggeration” concept from [22]. Here, we multiply all values of *p_i,j_* by a constant,γ, for the first 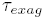 iterations. Because the *p_i,j_* values are large, this forces the *q_i,j_* values, which still must sum to one, to be as large as possible. This pushes the data’s natural clusters into tight, widely separated regions. For the work presented here, we use γ = 4 and 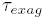 = 100. The stopping criteria for the algorithm is for the fractional improvement in the cost function to fall below 10^−4^.

#### 1. Selecting training-set points

Due to memory limitations (the memory requirements scale like *N*^2^), we can only embed a subsample of approximately 30,000 at one time. Although improving this number will be the subject of further research, our solution here is to generate an embedding based-upon a selection of 1,250 data points from each of the 24 individuals observed (out of ≈ 360,000 data points per individual). To ensure that these points create a representative sample, we perform t-SNE on 20,000 randomly-selected data points from each individual, using the same parameters as described above. This embedding is then used to estimate a probability density by convolving each point with a 2D gaussian whose whose width is equal to the distance from the point to its 10th nearest neighbor. This space is segmented by applying a watershed transform [23] to the inverse of the PDF, creating a set of regions. Finally, points are grouped by the region to which they belong and the number of points selected out of each region is proportional to the integral over the PDF in that region. This is performed for all data sets, yielding a total of 30,000 data points.

#### 2. Comparing new points to the training set

Given the embedding resulting from applying t-SNE to our training set, we embed additional points into our behavioral space by comparing each to the training set individually. Mathematically, let *X* be the set of all feature vectors in the training set, *Y* be their associated embeddings via t-SNE, *z* be a new feature vector that we would like to embed according to the mapping between *X* and *Y*, and *ζ* is the embedding of *z* that we would like to determine.

As with the t-SNE cost function, we will embed *z* by enforcing that its transition probabilities in the two spaces are as similar as possible. Like before, the transitions in the full space, *p_j|z_*, are given by

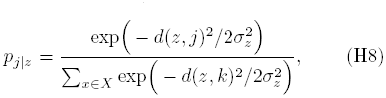

where *d*(*z, j*) is the Kullback-Leibler divergence between *z* and *x* ∈ *X*, and *σ_z_*is once again found by constraining the entropy of the condition transition probability distribution, using the same parameters as for the t-SNE embedding (2*^H_z_^* = 30 ± 10^−5^). Similarly, the transition probabilities in the embedded space are given by

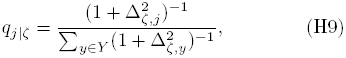

where 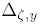 is the Euclidean distance between ζ and *y* ∈ *Y*.

For each *z*, we then seek the *ζ*\* that minimizes the Kullback-Leibler divergence between the transition probability distributions in the two spaces:

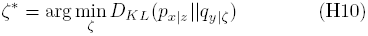

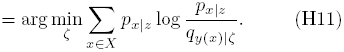

As before, this is a non-convex function, leading to potential complexities in performing our desired optimization. However, if we start a local optimization (using the Nelder-Mead Simplex algorithm [44, 45]) from a weighted average of points, *ζ*_0_, where

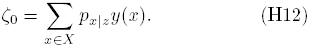

This point is usually within the basin of attraction of the minimum (see Figure 14) so that the global minimum is found. To ensure that this is true in all cases, we also evaluate the cost function on a coarse grid centered about *ζ*_0_ (25 × 25 evaluation points placed in a square lattice with a grid spacing of 1.25). If the cost function value was less than or within 10% of the previous minimum value, a new local optimization was run, starting from that location. This was necessary less than 1% of the time.

**FIG. 14.**
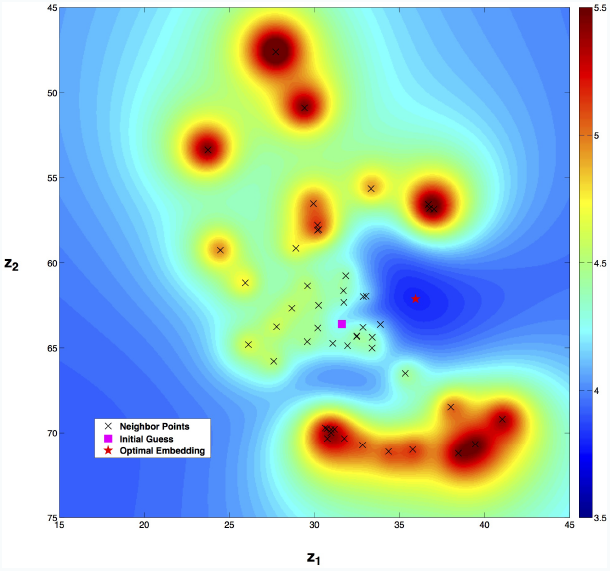
Cost function landscape for embedding a new point into the training set. The colormap represents the KL-divergences (Equation H10) at various points in the landscape. The black points are the nearest neighbor positions used in the calculation, the magenta square is the initial guess from Equation H12, and the red star is the found optimal embedding location.

Because this can be calculated independently for each value of *z*, the algorithm scales linearly with the number of points. We also make use of the fact that this algorithm is embarrassingly parallelizable. Moreover, because we have set our perplexity (2*^H^*) to be 30, there are rarely more than 50 points to which a given *z* has a non-zero transition probability. Accordingly, we can speed up our cost function evaluation considerably by only allowing *p_x|z_* > 0 for the nearest 200 points to *z* in the original space.

### Appendix I: Definition of behavioral states

#### 1. Resting and moving states

Observing the spatio-temporal dynamics in our behavioral space, it becomes apparent that movements can be described by relatively long spans of stability followed by quick bursts of movement (see Figure 17, blue line for a typical sequence). This can be quantified through the log-histogram of embedded-space speeds (Figure 3D), which displays a bimodal distribution. Decomposing this distribution into two log-normal distributions via expectation maximization [46], we now define the probability of a point to be in a pause state as the posterior probability that the data point in question belongs to the left-most distribution. The fly is said to be within a “behavioral state” at a moment in time if there is at least a .05 second (5 frame) sequence in the pause state, and the point in question does not cross over a region boundary (see following section). It is this definition that is used to compute transition probabilities such as those displayed in Figure 4A of the main text.

#### 2. Spatial segmentation

Given the strong clustering of the embedded space observed in Figure 3 of the main text, we would like to break this space into regions corresponding to these groupings. Many clustering algorithms have been developed for similar purposes [47], all with their respective strengths and weaknesses. Here, we choose a method that allows us to avoid pre-selecting the number of clusters by hand. Specifically, we create a composite image, *Β*_*σ*_, from our embedded data points 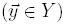, where,

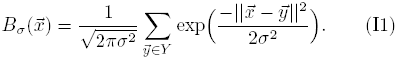

**FIG. 15.**
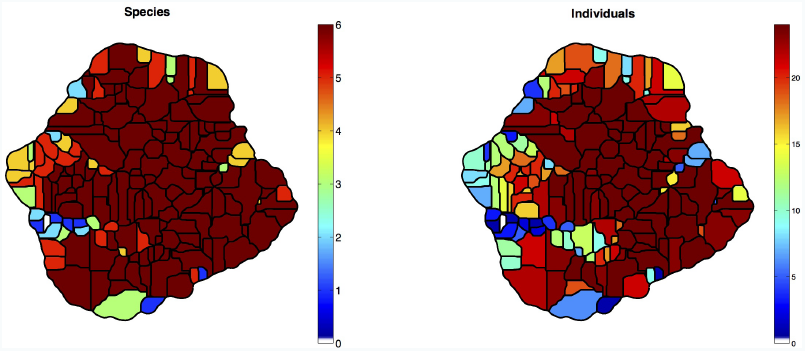
All flies visit most of behavioral space. The color coding here represents the number of species (left) and individual flies (right) that performed the stereotyped behavior corresponding to the region in the embedded space at some point during recording.

**FIG. 16.**
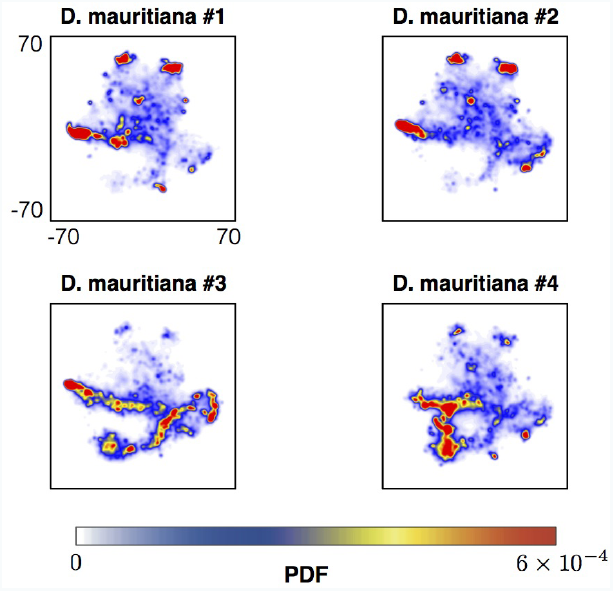
PDFs for each of the four *D. mauritiana* analyzed here, plotted separately.

**FIG. 17.**
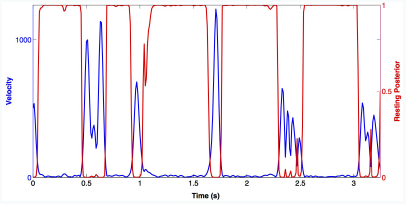
A typical dynamical sequence in the embedded space. The blue line shows quick bursts of velocity, followed by longer slow periods. The red line shows the posterior probability of belonging to the left-most distribution in Figure2D.

This image (as seen in Figure 2B of the main text) can be viewed as an estimate of the probability density function over the embedded space. Given this image, we can proceed to segment space into various regions, each associated with a local maximum of the PDF, via the watershed algorithm [23]. Additionally, we ignore regions of the space where *B_σ_* is sufficiently small (here, we ignore all areas where *B_σ_* < 5 × 10^−6^). Naturally, the number of regions found via this approach decreases monotonically with increasing *σ*. Accordingly, instead of specifying the number of clusters, we provide a minimum length scale beneath which all local anisotropies are smoothed. For the results presented in the main text, we choose *σ* = 1, as it provides a good balance between preserving local structure and elimination of spurious noise (*σ* is approximately equal to the length scale of spatial fluctuations seen when the trajectory remains in the “resting” state).

It should be noted, however, that the technique used here is the not the only method of partitioning the embedded points or space. For our purposes, however, it serves as an intuitive segmentation of space into natural clusters that is found to correspond well to stereotyped behaviors. Future work will include a focus on developing and applying a more formal approach to this clustering problem.

#### 3. Calculating transition fluxes

In order to perform our graph partitioning, we first must define a transition matrix, *T*, that will serve as our weighted graph. Here, we assume that *T_i,j_* is the relative transitional flux from behavior *i* to behavior *j*. In other words, the number of transitions observed from *i* to *j* divided by the total number of transitions observed altogether (after setting all self-transitions to zero). More specifically, if *P* (*i* → *j*) is the conditional probability that state *j* is the next state observed after observing state *i*, and *ρ_i_* is the fraction of the time that the system is observed in state *i*, then we can define our transition flux via

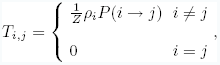

where

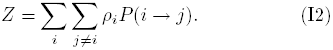

#### 4. Graph partitioning

To perform the graph partitioning of a given transition matrix, *T*, as displayed in Figure 3 and Figure 4 of the main text, we rely on the self-consistent Network Information Bottleneck (NIB) method introduced in [24]. This approach, taking its inspiration from the information bottleneck method [48], clusters a set of data points, *X*, into a group of modules, *Z*, in a manner such that the mutual information between *X* and *Z* is minimized (maximally compressing the data), given that the loss of mutual information between *X* and a third variable,*Y*, is constrained to be below a prescribed value. Stated more colloquially, one desires to pass the information that *X* preserves about *Y*, usually referred to as a ”relevance variable,” as reliably as possible despite having to pass through a bottleneck, *Z*.

This can be achieved through minimizing the functional, 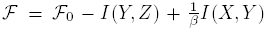 with respect to 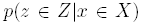. Here, 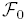 is a constant that does not depend on *p*(*z*|*x*) and *β* is a Lagrange multiplier that is monotonically related to amount of information loss allowed between *X* and *Y*. In practice, one solves this by iterating through the Blahut-Arimoto algorithm (see [24, 48] for details).

For our purposes, *X* is the set of all behavioral states obtained through the methods described in the previous section, *Z* is the set of modules, and *Y* is defined as the state at which one finds a random walker on *T^(sym)^* (the symmetrized version of *T*) after *τ* time steps. Here, we set *τ* to be the characteristic time scale of the transition matrix (the inverse of the second-largest eigenvalue of *T*). Accordingly, we can define *p*(*y*|*x*) ≡ exp(—*Lτ*), where *L* is the graph Laplacian of *T*^(*sym*)^, and *p*(*x*) is the steady-state probability distribution from *T*^(*sym*)^, which is proportional to the eigenvector corresponding to the matrix’s smallest non-zero eigenvalue. Starting with this information, a random initialization for *p*(*z*|*x*), and a pre-specified number of clusters, *k*, we solve the iterated equations for a relatively small value of *β*. We then anneal the system by taking the output from the previous iteration and starting a new iteration at a higher value of *β*, using the previous *p*(*z*|*x*) as an initial condition. Here, we start with *β* = 2 and anneal up to *β* = 1, 000, stopping each iterative algorithm with a relative convergence criterion of 10^−6^. The number of modules ranged from 2 to 12, for 20 replicates each, keeping the solution with the minimal value for 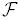.

**FIG. 18.**
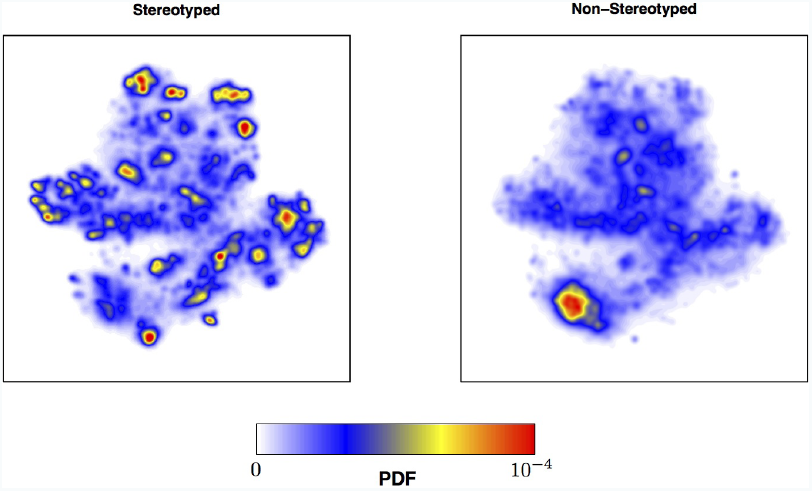
Probability density functions of positions in behavioral space conditional on belonging to a stereotyped behavior (Left), or not belonging to a stereotyped behavior (Right). Note the drastic smoothing out of the landscape in the nonstereotyped case.

**Table II.** Behavioral statistics for six species

| Species | # of States Visited | % of Time Stereotyped | Transitions / min |
| --- | --- | --- | --- |
| *D. mauritiana* | 155 | 51.8 | 104.2 |
| *D. melanogaster* | 134 | 57.7 | 92.4 |
| *D. santomea* | 143 | 48.2 | 111.7 |
| *D. sechellia* | 154 | 47.8 | 110.7 |
| *D. simulans* | 157 | 42.6 | 106.9 |
| *D. yakuba* | 137 | 51.7 | 91.0 |

This still leaves us with determining the number of modules in *Z*, however. To select this, we applied the Newman-Girvan modularity measure [49],

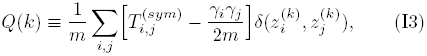

where 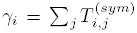,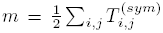 and 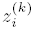 is the community assignment of state *i* given *k* modules in *Z*. We use the partition that corresponds to the maximal value of *Q(k)* (Figs. 20 & 21).

**FIG. 19.**
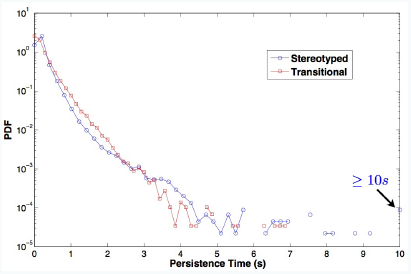
Histograms of persistence times for a fly remaining the stereotyped (blue) and transitional (red) states. Although rare, events as long as 15 seconds were found in the stereotyped state and are included within the last bin of the stereotyped histogram.

**FIG. 20.**
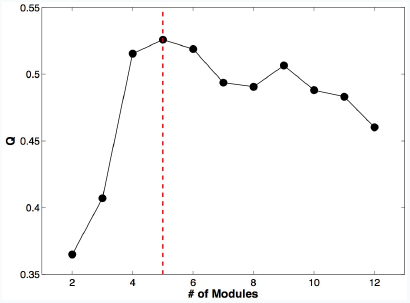
Modularity, *Q*, as a function of the number of included clusters for the *D. mauritiana*transition matrix.

**FIG. 21.**
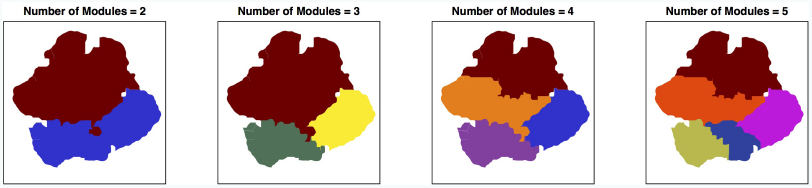
Graph partitions of the *D. mauritiana* transition matrix (Figure 3A of the main text) as a function of the number of modules included. Each color (arbitrarily ordered) represents a different module.

**FIG. 22.**
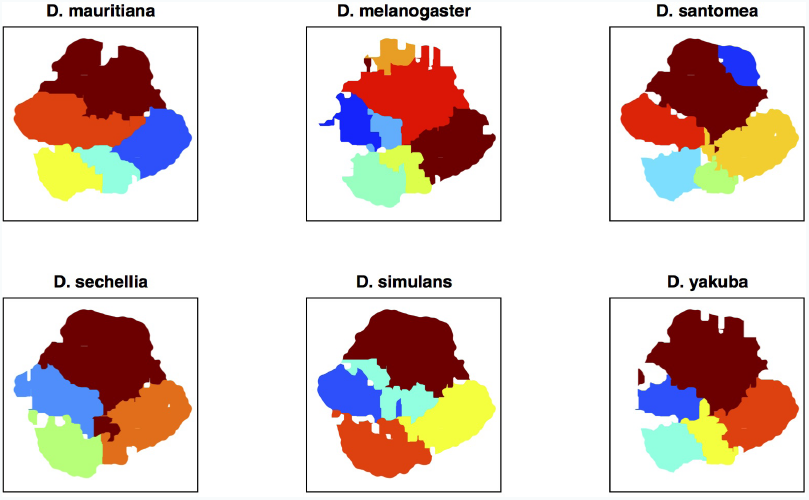
Graph partitions based on the transition fluxes for each of the six species. Each color (arbitrarily ordered) represents a different module. White holes represent states that were not observed in that particular species.

### Appendix J: Phase reconstruction and average orbits

The postural modes oscillate with a well-defined frequency for each non-resting stereotyped behavior (Fig. 24), implying, perhaps, that periodic changes in posture are mapped to points in behavioral space. This occurs even though the wavelet transform used to define the feature vectors does not preserve phase information. For example, in the locomotion sequences we observed, we find the postural eigenvalues oscillate regularly, with frequencies ranging from 2 to 14 Hz (Figs. 24 and 23). These frequencies are in good agreement with previous measurements of the fly walking and running gaits [50, 51].

**FIG. 23.**
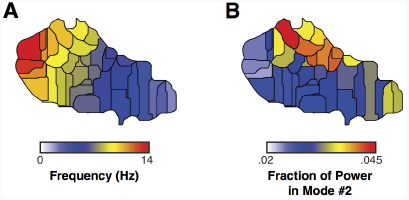
Map of dynamics within locomotion module. A) Map of the peak frequency of the wavelet transform as a function of position (averaged over all modes). Note that frequency monotonically decreases when going from left to right along the region. B) Map of the fraction of power in mode #2 for each region. Motion in mode #2 represents posterior movements (see main text, Fig. 1B). Here, the pattern goes from top to bottom.

**FIG. 24.**
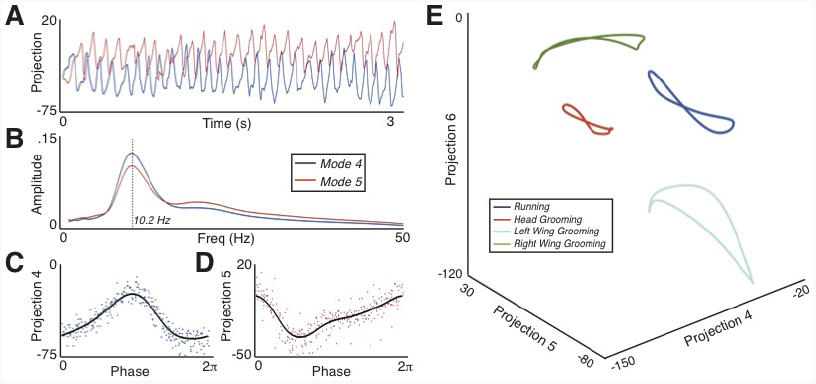
Behavioural space peaks are periodic orbits in postural space. (A) Projections onto the fourth (blue) and fifth (red) postural eigenmodes during a running sequence. (B) Average normalized wavelet transform value over this sequence for each of the two previous time series. (C) Fourth eigenmode projection values from (A) after being mapped onto the unit circle via phase reconstruction (dots) and the average curve using Von Mises weighted averaging (line). (D) Same as (C) but using the fifth eigenmode. (E) Plots of the phase-averaged curves for four different behavioral sequences. The inset displays the respective regions of space associated with each of the curves (same color code).

Mapping these projections onto the unit circle using a phase reconstruction algorithm [18], we construct an average path through postural space (Fig. 24C–D).

For phase reconstruction of periodic orbits, we use the Phaser algorithm originally introduced in [18]. Phaser takes a set of synchronized oscillators and uses the measurements of these values to map measurement times onto the unit circle. This algorithm works by using Hilbert transforms to construct phase estimations from each oscillator separately, then recombining to form a maximum likelihood estimate utilizing Fourier-series based corrections. Here, our coupled oscillators are the postural eigenmode projections, *y_k_* (t). Accordingly, the returned phase is a combined estimate from observing all 50 modes at once. Given at least 7 oscillatory cycles, this method provides a robust phase estimation in the limit where dynamics are deterministic with only small variations with respect to the mean trajectory.

We find oscillatory postural dynamics for other stereotyped behaviors, with each behavior resulting in a periodic orbit in postural space (Fig. 24E). Periodic orbits are suggestive of underlying low-dimensional dynamic attractors that produce stable behavioral templates. These types of motifs have been hypothesized to form the basis for neural and mechanical control of legged locomotion at fast time scales [4, 6]. The large number of behavioral bouts that we observe for each data set will allow future data-driven analyses of the orbit stabilities and their dominant slow degrees of freedom[18, 52].

From these phase reconstructions, we find the average orbit through a von Mises distribution weighted average. More precisely, we construct the average orbit for eigenmode *k*, *μ*^(*k*)^ (*ϕ*) via

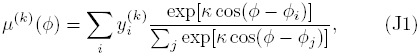

where 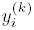 is the projection onto the *k^th^*eigenmode at time point *t_i_*, *ϕ_i_*is the phase associated with the same time point, and *κ* is related to the standard deviation of the von Mises distribution (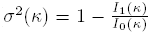, where *I_v_*(*x*) is the modified Bessel function of *ν*^*th*^ order). Here we find the value of *κ* ≈ 50.3, which is the *κ* resulting in *σ* = .1.

### Appendix K: Calculating JS-divergences

To calculate the distance between our behavioral probability distributions (Fig. 4 of the main text), we rely on the Jensen-Shannon divergence [25], defined via

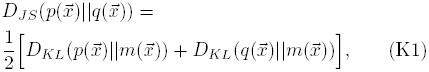

where 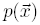 and 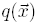 are probability distributions, *D_KL_* is the Kullback-Leibler divergence and

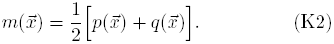

This quantity (bounded by zero and one) can be interpreted as the mutual information between draws out of the mean distribution, *m*, and a binary string that is 1 if the draw is from *p* and 0 if from *q*. In other words, this is a measure of distinguishability. If *D**_JS_**(p||q)* = 1, this means that given a draw from *m*, it is possible to precisely determine which of *p* or *q* it came from. As the divergence approaches zero, the number of draws needed to determine which distribution one is drawing from increases monotonically.

### Appendix L: Supplementary Movies

#### Movies available upon request at

**Movie 1.** Raw video data of a behaving fly (left) and the corresponding segmented and aligned data (right).

**Movie 2.** Dynamics in behavioral space. Raw video of a behaving *D. mauritiana* (middle) is displayed alongside coordinates of the fly’s position within the filming apparatus (left) and its position in the embedded behavioral space (right). The red circles represent the positions in the appropriate coordinate system and the trailing lines are the positions traversed in the previous .5 s. The light blue shading indicates that a particular behavior is being performed, and the blue text below the video of the fly gives a coarse label for the behavior. The first portion of the movie is 5 s, played at real time (indicated by “Real Time” above the fly video), and the subsequent portion of the movie is slowed down by a factor of 6 for clarity (indicated by “Slowed 6×”).

**Movies 3-10.** Each movie is a mosaic of multiple instances of specific regions in behavioral space as displayed in Fig. 25 and Table III. Every movie contains multiple segments from each of the six examined species (four examples per species). All films are slowed by a factor of 4 for clarity.

**Table III.** Supplementary Movies

| Movie | Label |
| --- | --- |
| 3 | Left wing grooming |
| 4 | Wing flicking |
| 5 | Right wing grooming |
| 6 | Right leg flicking |
| 7 | Head grooming |
| 8 | Walking (non-tripod gaits) |
| 9 | Front legs grooming |
| 10 | Proboscis extension |

**FIG. 25.**
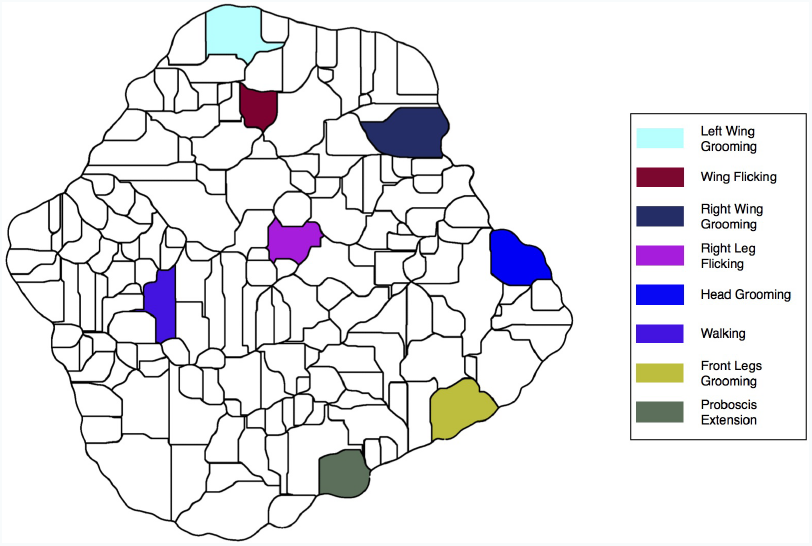
Stereotyped behaviors shown in Movies 3 to 10. Black lines are the boundaries resulting from the watershed segmentation described in Section I2. Colored regions are the areas of behavioral space corresponding to the movie

**Movies 11.** Comparison between typical running gaits for the three species of the *D. simulans* clade. Top: composite movies for *D. mauritiana, D. sechelia*, and *D. simulans* (slowed by a factor of 4). For each species, the state chosen is the region containing the peak of the PDF within the locomotion region (Fig. 4E). Note the relatively fast gait for *mauritiana*, the slower gait seen in *simulans*, and the subtle wing motions occurring during the gaits of *sechellia.* Bottom: the region from which the gaits are shown from each species.

